# Data-driven inference of crosstalk in the tumor microenvironment

**DOI:** 10.1101/835512

**Authors:** Umesh Ghoshdastider, Marjan Mojtabavi Naeini, Neha Rohatgi, Egor Revkov, Angeline Wong, Sundar Solai, Tin Trung Nguyen, Joe Yeong, Jabed Iqbal, Puay Hoon Tan, Balram Chowbay, Ramanuj DasGupta, Anders Jacobsen Skanderup

## Abstract

Signaling between cancer and nonmalignant (stromal) cells in the tumor microenvironment (TME) is key to tumorigenesis yet challenging to decipher from tumor transcriptomes. Here, we report an unbiased, data-driven approach to deconvolute bulk tumor transcriptomes and predict crosstalk between ligands and receptors on cancer and stromal cells in the TME of 20 solid tumor types. Our approach recovers known transcriptional hallmarks of cancer and stromal cells and is concordant with single-cell and immunohistochemistry data, underlining its robustness. Pan-cancer analysis reveals previously unrecognized features of cancer-stromal crosstalk. We find that autocrine cancer cell cross-talk varied between tissues but often converged on known cancer signaling pathways. In contrast, many stromal cross-talk interactions were highly conserved across tumor types. Interestingly, the immune checkpoint ligand PD-L1 was overexpressed in stromal rather than cancer cells across all tumor types. Moreover, we predicted and experimentally validated aberrant ligand and receptor expression in cancer cells of basal and luminal breast cancer, respectively. Collectively, our findings validate a data-driven method for tumor transcriptome deconvolution and establishes a new resource for hypothesis generation and downstream functional interrogation of the TME in tumorigenesis and disease progression.

---

The tumor microenvironment (TME) is a multi-faceted cellular environment that both constrains the evolving tumor (Hanahan and Coussens, 2012) and plays a pivotal role in tumor progression and therapeutic response (Junttila and de Sauvage, 2013). Existing experimental methods to characterize the TME, such as imaging and single-cell based approaches, cannot be applied retrospectively to existing large-scale bulk tumor datasets, representing a vast and mostly unexplored resource for studying cancer cell ligand-receptor repertoires and cross-talk in the. Existing approaches to deconvolve bulk tumor gene expression profiles have mainly focused on estimation of cell type fractions (Aran et al., 2017; Newman et al., 2015) or have been optimized or tested on a limited set of tumor types (Ahn et al. 2013; Moffitt et al., 2015; Quon et al., 2013; Wang et al., 2018).

Previous studies have analysed tumor purity in a pan-cancer context (Aran et al., 2015) and developed regression-based methods for accurate mixed-tissue transcriptome deconvolution (Shen-Orr et al., 2010). Here we combined these approaches to estimate cancer and stromal (comprising any non-cancer cell) compartment molecular profiles from bulk tumor RNA-seq data and infer signaling interactions between average representative cells in these two compartments. Uniquely, our approach (TUMERIC) avoids making assumptions about the transcriptional profiles of cancer and stromal cells in a given tumor by integrating independent estimates of tumor purity derived from DNA sequencing and copy number profiling data. We validate the approach using public cancer genomics data, mass spectrometry protein abundance data, single cell RNA-seq data, and immunohistochemistry imaging data. We apply the method to ∼8000 tumor samples across 20 solid tumor types from The Cancer Genome Atlas (TCGA) to infer cancer and stromal cell crosstalk conserved across tumor types as well as tissue specific interactions. Finally, we compare cross talk across the molecular subtypes of breast cancer and infer signaling interactions specific to the aggressive basal subtype.

## Results

### Overview of approach

We first estimate the cancer cell fraction (tumor purity) from somatic mutation allele frequencies, DNA copy number profiles, and mRNA expression signatures of the tumors using a robust consensus approach (Fig. 1 and Methods). The tumor transcriptome profiles are then deconvolved into an average cancer and stromal cell profile using non-negative linear regression. To infer cancer and stromal cell ligand and receptor repertoires, as well as potential crosstalk between these compartments, we combine the inferred expression profiles with a curated database of ligand-receptor (LR) interactions (Ramilowski et al., 2015) to estimate relative LR complex concentrations under equilibrium (Methods).

**Figure 1:**
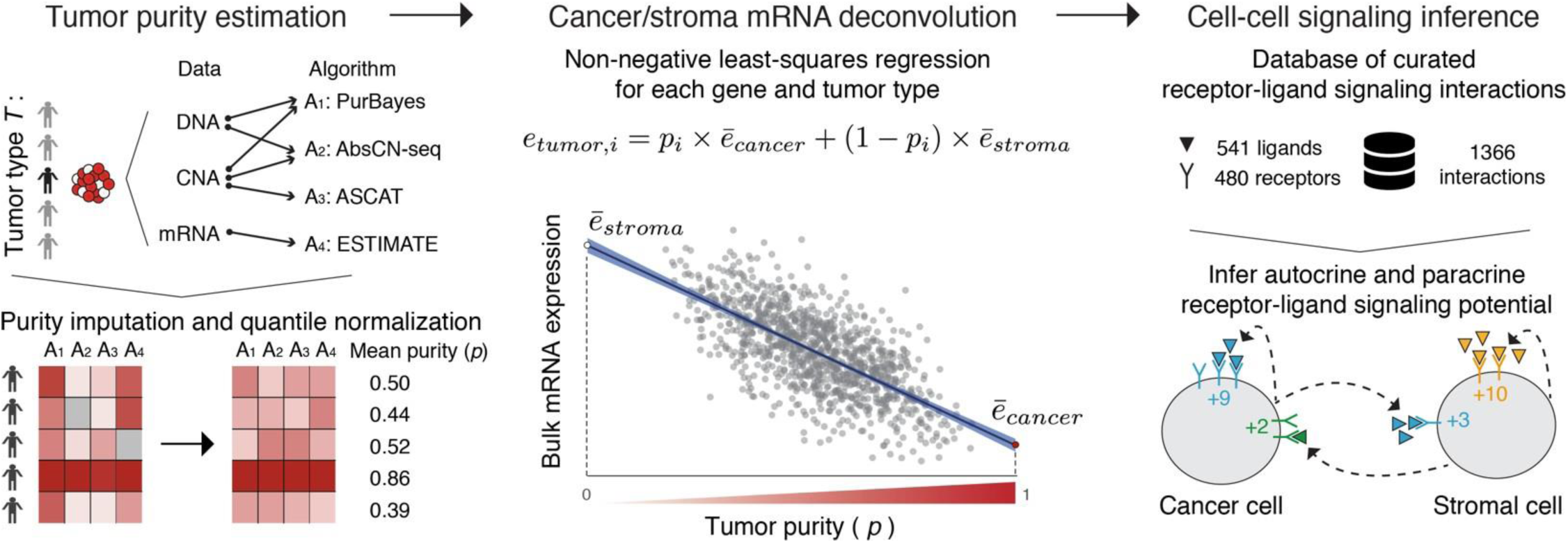
Overview of approach. The purity of each bulk tumor sample is first estimated using a consensus approach. mRNA expression levels in “average” cancer and stromal cells are inferred for a set of tumors (e.g. tumor type) using non-negative least-squares regression; figure shows data for *CD4* in breast cancer. Candidate autocrine and paracrine signaling interactions are inferred using a database of curated receptor-ligand signaling interactions.

### Estimation of tumor purity across 8000 tumor samples

To analyze crosstalk in a pan-cancer context, we first estimated tumor purities of ∼8000 samples across 20 solid tumor types from TCGA. Briefly, we obtained purity estimates based on DNA somatic variant allele frequency (PurBayes and AbsCN-seq), copy number (ASCAT), and mRNA expression (ESTIMATE) data using four existing algorithms, followed by imputation and quantile normalization, to produce robust consensus tumor purity estimates across all sample. Most tumors had a purity in the range 40-70%, but there was large variation within and between tumor types (Fig. 2a, Suppl. Fig. 1). Pancreatic adenocarcinoma (PAAD) tumors had very low purity (median ∼39%), consistent with previous observations (Wood and Hruban, 2012). The glioblastoma (GBM) and ovarian cancer (OV) samples had the highest purity estimates, likely influenced by sample selection bias in the first phase of the TCGA project.

**Figure 2:**
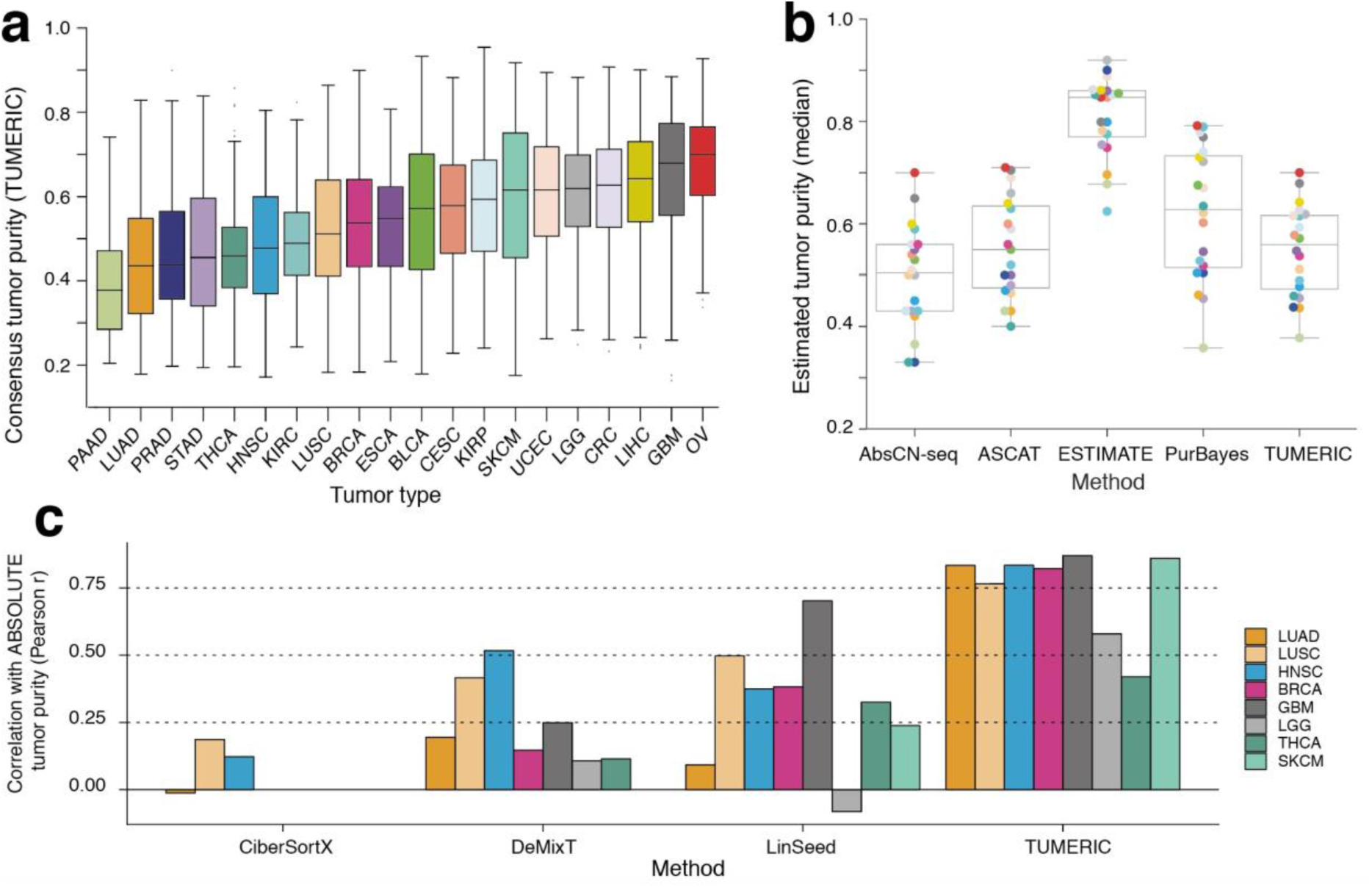
Consensus tumor purity estimation. **a**) TUMERIC consensus tumor purity estimates for ∼8000 bulk tumor samples across 20 solid tumor types. **b**) Median tumor purity of each tumor type as estimated by 4 different methods, including TUMERIC consensus estimates. Points (representing median purity across tumor type) are color coded according to tumor types in panel a). **c**) Concordance of tumor purity estimates from existing transcriptome deconvolution methods with ABSOLUTE (DNA-derived) tumor purity estimates; TUMERIC consensus purity estimates included for comparison.

We found that previously published consensus tumor purity estimates (Aran et al., 2015), while positively correlated with TUMERIC estimates, were likely overestimating purity by >30% as compared to genome-derived purity estimates (Suppl. Fig. 1). We then compared TUMERIC consensus purity values with purity estimates from recently published transcriptome deconvolution methods, using tumor purity estimates computed by the TCGA pan-cancer consortium (ABSOLUTE method) as an independent benchmark (Hoadley et al., 2018). One method (CiberSortX) could only be applied in three tumor types because it required a tumor type specific signature matrix for deconvolution. Another method (DeMixT) could only be applied in tumor types where matched normal tissue samples were available. Overall, the concordance with ABSOLUTE purity was generally low for the three tested transcriptome deconvolution methods (Pearson *r* > 0.4 in only 2/8 tested tumor types, Figure 2c). Furthermore, all methods failed (*r* < 0.15) to estimate tumor purity in at least two of the tested tumor types. In contrast, TUMERIC consensus purity estimates were generally highly correlated (*r* > 0.75 in 6/8 tumor types, lowest *r* = 0.42 for THCA) with TCGA ABSOLUTE purity estimates across cancer types.

### Validation of approach

We performed multiple analyses to evaluate the accuracy of TUMERIC in deconvolution of cancer and stromal compartment transcriptomes. Firstly, known stromal (*FAP, CD3D, CD4, CSF1R*) and epithelial-cell specific factors (*EPCAM*) showed expected strong and consistent gene expression differences between stromal and cancer compartments across cancer types (Suppl. Fig. 2-6). Next, since somatic copy number alterations (CNAs) are hallmarks of cancer cell genomes, we reasoned that genes expressed exclusively in the stromal compartment should not be affected by tumor CNAs. We used TUMERIC to infer the top-100 cancer and stromal-cell specific genes in each tumor type. The cancer-specific genes in each tumor type were distributed widely across the genome (Suppl. Fig. 17). We found strong correlation between bulk tumor CNA and expression of cancer-specific genes, but no correlation between tumor CNA and expression of stroma-specific genes (Fig. 3a). Variation in correlation between tumor types could be explained by the overall prevalence of CNAs in a given tumor type, where tumor types with higher levels of chromosomal instability showed higher correlation of tumor CNA and expression of cancer-specific genes (Fig. 3a). Similarly, as a positive control, we found that previously derived stromal and immune-cell specific genes (Yoshihara et al., 2013) were inferred by TUMERIC to have markedly higher expression in the stroma compartment of all tumor types (Fig 3b).

**Figure 3:**
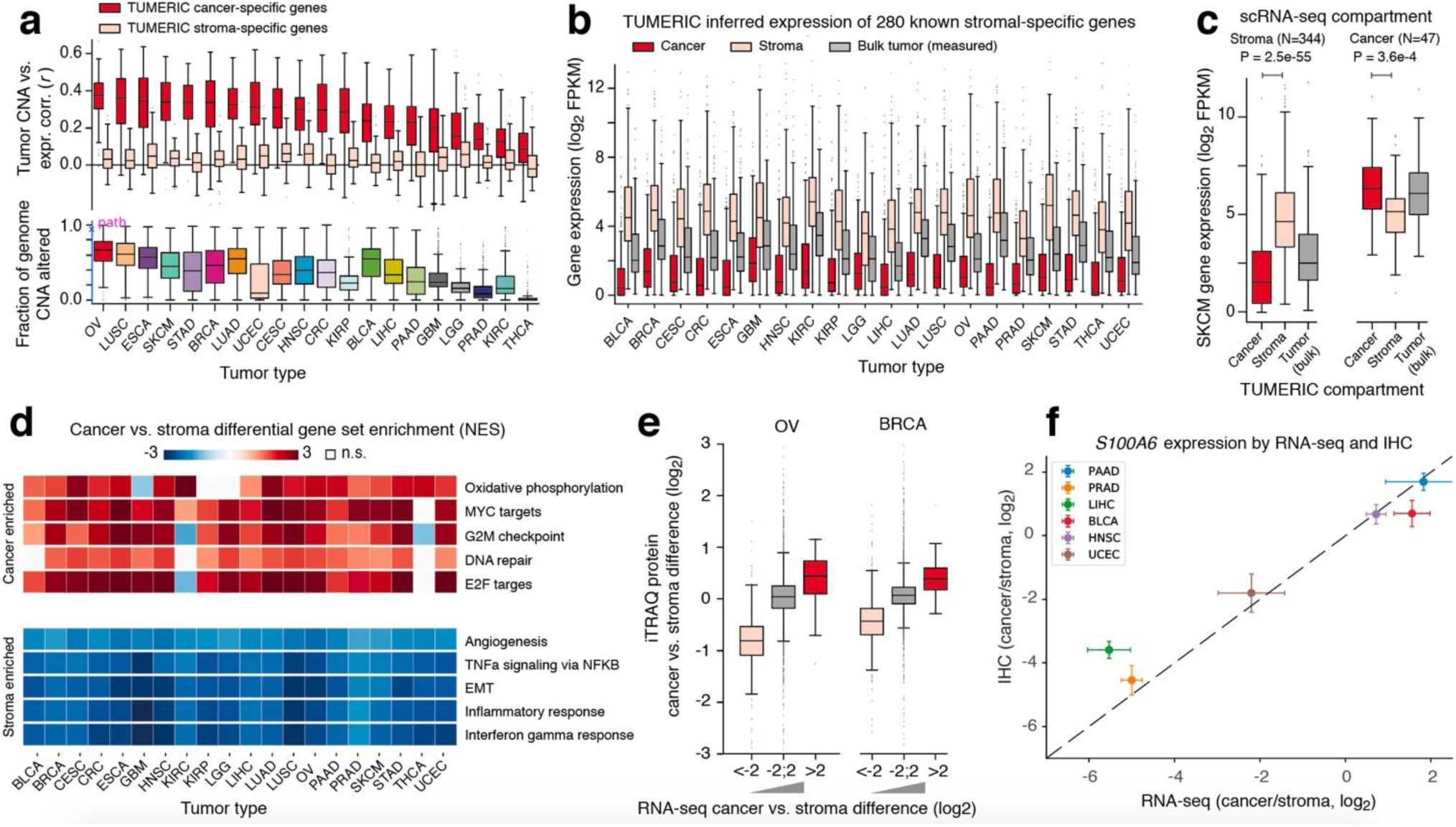
Validation of tumor gene expression deconvolution. **a**) Genes specifically expressed in cancer and stromal cells were inferred for each tumor type, and the correlation (Pearson *r*) between mRNA expression and somatic copy number alterations (CNA) at each gene locus was evaluated (top panel). Tumor types are ordered by the difference in means of cancer and stromal gene correlations. The fraction of genome altered by CNA was determined for each tumor sample (bottom panel). **b**) Inferred cancer and stroma compartment expression levels for 280 known stromal-specific genes. **c**) Inferred cancer and stroma compartment expression levels in melanoma (SKCM) for cancer and stroma specific genes previously identified with melanoma scRNA-seq. **d**) Genes were ordered by inferred expression difference between cancer and stroma compartments in each tumor type, and gene set enrichment analysis (GSEA) was used to identify cancer and stroma enriched gene sets. **e**) Protein expression was inferred for cancer and stroma compartments in (OV) and breast (BRCA) cancer cohorts using iTRAQ protein quantification data and compared to deconvolved RNA-seq expression data. **f**) Comparison of immunohistochemistry (IHC) and deconvolved RNA-seq expression for a gene (*S100A6*) with highly variable cancer/stroma expression across cancer types; error bars reflect standard deviations of the point estimates.

To test the concordance of TUMERIC with tumor single-cell RNA-seq (scRNA-seq) profiling, we analyzed TUMERIC expression estimates for cancer and stromal-cell specific genes identified by scRNA-seq of melanoma tumors (Tirosh et al., 2016). TUMERIC inferred significantly higher stroma-compartment expression for scRNA-seq stromal-cell specific genes (P=2e-55, Mann Whitney, two-tailed), and significantly higher cancer-compartment expression for cancer-cell specific genes (P=3.6e-4, Fig. 3c). Next, using gene set enrichment analysis, we found consistent association of compartment-wise inferred gene expression and known hallmarks of cancer (e.g. cell cycle and DNA repair) and stromal cells (e.g. angiogenesis and immune response) across cancer types (Fig. 3d, Suppl. Fig. 7). We then evaluated the extent that the deconvolved mRNA profiles represent an accurate proxy for protein levels in the cancer and stromal cells. We used TUMERIC to deconvolve protein abundance data from two TCGA tumor types (Edwards et al., 2015), and found strong concordance between the mRNA and protein expression profiles (Fig 3e). Finally, to test the concordance of TUMERIC with immunohistochemistry (IHC) data, we identified genes with high variability of cancer and stromal cell differential expression across tumor types and used IHC data from The Human Protein Atlas (Uhlen et al., 2017) to confirm that expression patterns of two such genes were indeed variable and concordant across tumor types (Fig 3f and Suppl. Fig 8). Although this IHC comparison is limited to two genes, the combined concordance of transcriptome deconvolution data with both proteomic and IHC data supports an overall concordance between transcriptome and protein-level data. Overall these diverse comparisons with orthogonal types of data support that TUMERIC can robustly deconvolve gene expression profiles of cancer and stromal cell compartments across the 20 tested tumor types.

### Compartment specific expression of ligands and receptors

To explore ligand-receptor (LR) signaling across cancer types, we focused on 263 ligands and 242 receptors (603 LR pairs) with detectable bulk tumor gene expression (>1 RPKM in >25% of samples) in at least half of the 20 cancer types. We first analyzed for compartment specificity of these ligands and receptors. Expectedly, ligands belonging to the complement system (e.g. *C1QB* and *C3*) as well as leukocyte specific chemokines (e.g. *CCL5* and *CCL21*) had high stroma specific expression across tumor types (Fig 4a). Cancer-specific ligand expression across tumor types was much less frequent and pronounced. Among the most common cancer-specific ligands were endothelial cell targeting factors (*PODXL2* and *VEGFA*), consistent with the hypothesis that cancer cells interact with and induce tumor vascularization. Immune (e.g. *CD2* and *CD4*) and macrophage cell (*CSF1R*) specific factors were expectedly among the top stromal specific receptors across tumor types (Fig 4b). Similar to ligands, receptors with cancer-cell specific expression across tumor types were less common. The top common cancer specific receptors included known cancer associated members of the EGF-family (*ERBB2* and *ERBB3*), Wnt-family (*LRP4, LRP5* and *LRP6*), and FGF-family (*FGFR3*). In summary, these data underline that our approach is able to tease apart cancer and stromal-compartment specific expression of ligands and receptors from bulk tumor transcriptome profiles.

**Figure 4:**
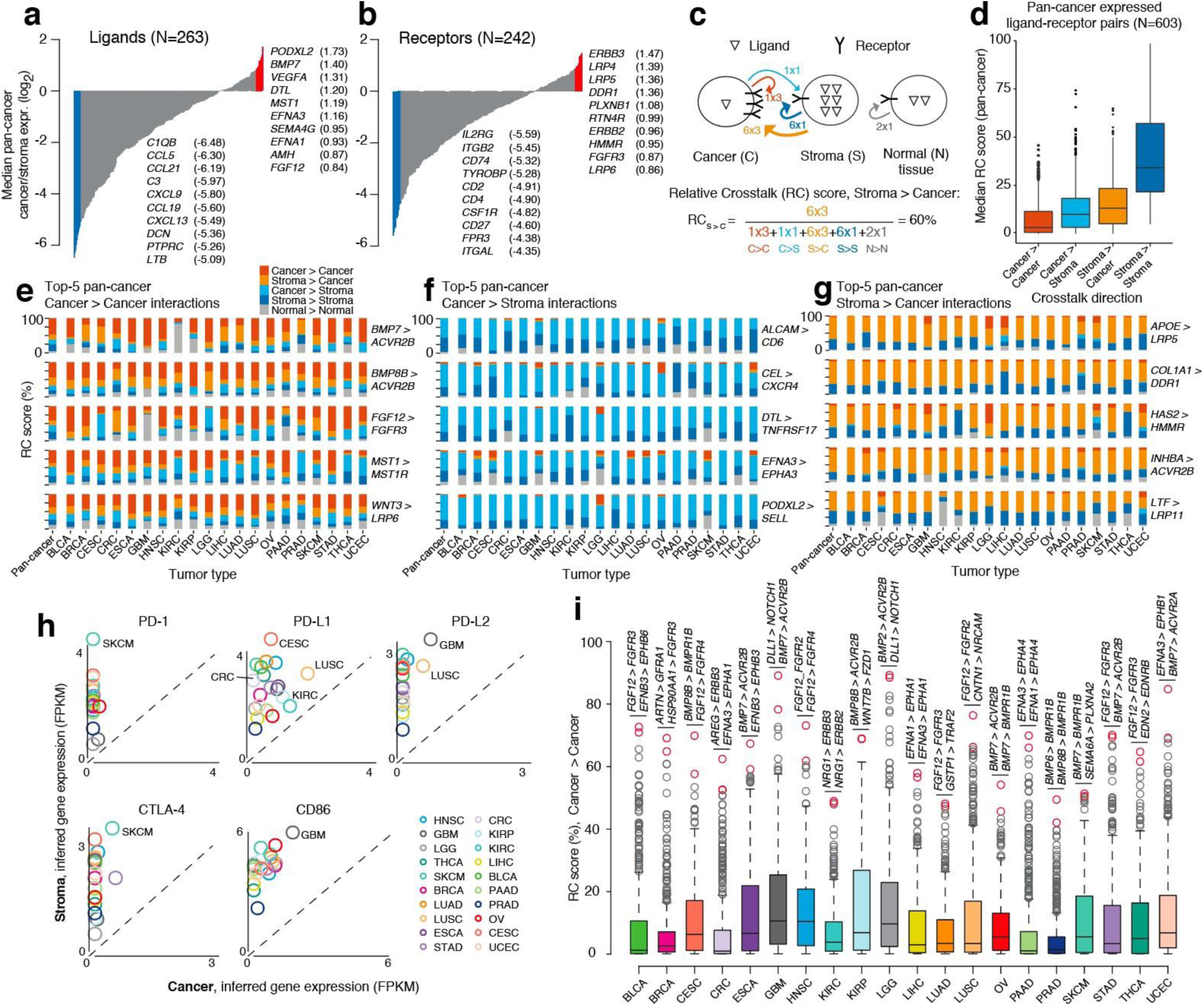
Pan-cancer inference of crosstalk. **a, b**) Differential expression (median) between cancer and stroma compartments of ligands **a**) and receptors **b**) across cancer types. **c**) The Relative Crosstalk (RC) score approximates the relative flow of signaling in the four possible directions between cancer and stromal cell compartments, including a bulk (non-deconvolved) matched normal tissue reference. In the concrete example, a given ligand is expressed with 1, 6, and 2 copies (receptor in 3, 1, and 1 copies) in cancer, stroma, normal cells, respectively. This yields a stroma-to-cancer (S>C) RC score of 60%. **d**) Median RC score across the 20 solid tumor types was estimated and plotted for each direction of signaling. **e, f, g**) RC scores for the 5 ligand-receptor pairs with highest median pan-cancer cancer-cancer autocrine **e**), cancer-stroma **f**), and stroma-cancer **g**) paracrine signaling scores. **h**) Inferred cancer and stromal compartment gene expression across tumor types for immune checkpoint ligands (PD-L1, PD-L2, CD86) and receptors (PD-1, CTLA-4). **i**) Distributions of RC scores for cancer-cancer autocrine signaling across tumor types; the top-2 ligand-receptor pairs are highlighted for each tumor type.

### Recurrence of ligand-receptor interactions across tumor types

Next, we developed the Relative Crosstalk (RC) score to quantify and differentiate between types of autocrine and paracrine (here denoting signaling within or between compartments, respectively) ligand-receptor (LR) crosstalk (Fig 4c and Suppl. Table 1). We first evaluated the extent that known LR pairs had consistent crosstalk directionality across tumors types and found a striking difference between the cancer and stromal compartment. While only 3 LR pairs had strong autocrine cancer signaling scores (median RC score > 40%) across tumor types, 264 LR pairs showed strong autocrine stroma signaling RC scores across cancer types (Fig 4d). This suggests that, in solid tumors, autocrine cancer signaling tends to be tumor type specific and determined by the cancer cell-of-origin. In contrast, stromal autocrine signaling is often conserved and independent of tumor type and tissue. Interestingly, the paracrine signaling interface between cancer and stromal cell compartments also had a high number of recurrent interactions (26-40 conserved interactions, respectively), highlighting the importance of the tumor environment on cancer cell biology.

Inferred recurrent cancer-cell autocrine LR pairs included receptors such as *FGFR8, LRP6* and *MST1R* (Fig. 4e). Signaling through *ACVR2B* was notably both among the top inferred cancer autocrine and stroma-to-cancer signaling interactions across tumor types (Fig 4e,g), highlighting a common role for ACVR-coupled TGF-beta/SMAD signaling in tumorigenesis. We analyzed recurrent cancer-to-stroma LR interactions to investigate how cancer cells may engage and shape their microenvironment (Fig 4f). Top inferred cancer-to-stroma interactions included the lymphocyte-specific selectin (*PODXL2*-*SELL*), highlighting a possible conserved interaction between cancer cells and leukocytes or endothelial cells (Fieger et al., 2003) (only the leukocyte-specific selectin (*SELL*) was abundantly expressed across most tumors and included in our analysis).

### PD-1 and CTLA-4 immune checkpoint ligands are overexpressed in stromal cells

The top recurrent stroma-stroma interactions included many chemokine signaling interactions, especially involving *CXCR3* and *CXCR5* receptors, suggesting common stroma induced recruitment of leukocytes and lymphocytes to solid tumors (Suppl. Fig 9). Interestingly, the top recurrent stromal interactions also included the known immune checkpoint interactions CD86/CTLA-4 (*CD86*/*CTLA4*) and PD-L2/PD-1 (*PDCD1LG2*/*PDCD1*). The known PD-L1/PD-1 (*CD274*/*PDCD1*) checkpoint interaction was not among our initial curated and analyzed ligand-receptor interactions, but a follow-up analysis of this LR pair revealed a similar stroma-stroma enriched cross-talk pattern (median stroma-stroma RC=87%). Interestingly, expression levels of immune checkpoint ligands (PD-L1, PD-L2, CD86) were highest in stroma across all tumor types. This suggests that the bulk of immune checkpoint inhibitory signals in the tumor microenvironment may be mediated by non-cancer cells. However, some tumor types showed moderate to high expression of especially PD-L1 and CD86 in cancer cells. Glioblastoma (GBM) was the tumor type with highest expression of both CD86 and PD-L2 in both stroma and cancer cells. In contrast, brain tumors (GBM and LGG) also had the lowest stromal expression of both the CTLA-4 and PD-1 receptor as compared to all other tumor types, highlighting potential differences in T-cell infiltration levels, activation states, or dynamics of checkpoint inhibition in brain tumor tissue.

### Expression of immune checkpoints and response to immune checkpoint inhibition

Expectedly, inferred gene expression of the two checkpoint receptors, PD-1 and CTLA-4, was high in stroma and almost absent in cancer cells (Fig. 4h). Stromal expression of these two receptors was highest in melanoma (SKCM), the solid tumor type where immune checkpoint blockade was first clinically approved and where therapeutic blockade currently shows the most profound response rates (Sharma and Allison, 2015; Yarchoan et al., 2017). PD-L1 was inferred to have high expression in cancer cells of cervical squamous cell carcinoma (CESC), lung squamous cell carcinoma (LUSC), and renal cell carcinoma (KIRC, KIRP) (Fig 4h, Suppl. Fig 9). These tumor types all have high response rates to PD-L1/PD-1 checkpoint inhibition (Yarchoan et al., 2017). In contrast, colorectal cancer (CRC) has one of the poorest response rates to checkpoint inhibition (Yarchoan et al., 2017) and also showed the lowest (near-zero) inferred cancer cell expression of PD-L1 as compared to other tumor types. Overall, these results indicate that bulk tumor deconvolution of immune checkpoint cell-cell interactions in cancer and stroma compartments may provide insights into deciphering the conditions for effective immune checkpoint therapy.

### Convergence of autocrine cancer cell crosstalk across tumor types

The pan-cancer analysis indicated that cancer-cell directed LR interactions tended to differ across tumor types. We therefore searched for LR interactions with extreme autocrine RC scores in individual tumor types. Indeed, we found that most (14/20) cancer types had multiple LR interactions with high cancer autocrine scores (RC score > 60%, Fig 4i). Interestingly, the top autocrine cancer-cancer LR interactions tended to converge on the same signaling pathways. 9/20 solid tumor types had (among top-2, Fig 4i) autocrine interactions converging on TGF-beta and SMAD signaling pathways (BMP and ACVR-receptors). 8 tumor types had interactions involving different fibroblast growth factor ligands and receptors, and 6 tumor types had interactions converging on Eph/ephrin signaling. These results suggest that cancer cells of solid tumors may often be dependent on autocrine activation of these signaling pathways, and our data highlight specific LR candidate interactions associated with this activation in each tumor type.

### Validation of compartment-specific ligand-receptor expression using scRNA-seq data

We next investigated compartment-specific expression of candidate ligand-receptor pairs using a melanoma scRNA-seq dataset (Tirosh et al., 2016). Firstly, we analyzed the 3 pan-cancer cancer-to-cancer pairs with highest inferred cancer-cancer signalling in SKCM (*BMP7* > *ACVR2B, BMP8B* > *ACVR2B, WNT3* > *LRP6*, Fig 4e). Among these pairs, the two receptors (*ACVR2B* and *LRP6*) were inferred to be overexpressed in cancer cells (∼4-fold higher expression) and ligands were inferred to have unaltered expression in SKCM (Suppl. Table 1). We then compared the expression of the 2 receptors in malignant and non-malignant cells as inferred by the authors (Tirosh et al., 2016). Confirming our results, both receptors were significantly overexpressed in the cancer compartment (*P* <1e-10, Wilcoxon rank-sum, Suppl. Fig. 16). Next, we analysed the top 2 inferred SKCM cancer autocrine pairs (*BMP7* > *BMPR1B, SEMA6A* > *PLXNA2*, Fig. 4i). Within each of these two pairs, the receptor *BMPR1B* and the ligand *SEMA6A* were inferred to be significantly overexpressed in cancer cells (4 and 7-fold, respectively, Suppl. Table 1). Confirming our results, both of these genes were also significantly overexpressed in cancer cells of the melanoma RNA-seq dataset (P <1e-10, Suppl. Fig. 16).

### Autocrine and paracrine crosstalk in brain tumors

The two brain tumor types (LGG and GBM) were both characterized by very high autocrine Delta-Notch signaling scores (*DLL1*-*NOTCH1*, RC > 80%, Fig 4i). Both *DLL1* and *NOTCH1* were inferred to have >4 fold higher expression in cancer compared to tumor stromal cells and normal brain tissue in both tumor types. *DLL1* cancer cell expression was ∼16-fold higher than stroma and normal tissue in LGG (Suppl. Fig 10). These data are consistent with previous studies showing that Notch autocrine/juxtacrine signaling is critical for glioma tumorigenesis (Purow et al., 2005; Teodorczyk and Schmidt, 2015).

In contrast to autocrine cancer cell signaling, stroma-to-cancer crosstalk is lost when cancer cells are cultured *in vitro*. Interestingly, the top-2 stroma-to-cancer specific interactions for glioblastoma (GBM) converged on *EGFR* (Suppl. Fig 11). *EGFR* is a well-studied oncogene in GBM where 40-50% of patient tumors have *EGFR* overexpression driven by gene amplification (Brennan et al., 2013). We inferred >30-fold higher *EGFR* expression in GBM cancer cells compared to the stromal compartment and normal brain tissue (Suppl. Fig 11). In contrast, all canonical EGFR ligands were expressed at highest levels in the stroma (Suppl. Fig 11). *SPINK1* and *AREG* ligands were inferred to be exclusively expressed in stroma, and the most abundant ligand, *EFEMP1*, had ∼8-fold higher expression in the stromal compartment compared to cancer cells and normal brain tissue. A similar pattern was observed in lower grade gliomas (LGG) (Suppl. Fig 11), suggesting that EGFR signaling in glioma and glioblastoma cancer cells is driven mainly by ligands produced by the tumor infiltrating stroma. These data are also consistent with the observation that glioblastoma cancer cells rapidly loose EGFR amplifications during cell culture (Pandita et al., 2004; William et al., 2017).

### Crosstalk associated with subtypes of breast cancer

To investigate whether TUMERIC could identify differences in crosstalk between subtypes of a given tumor type, we inferred cross-talk associated with basal-like triple-negative breast cancers (TNBCs), a more aggressive subtype of breast cancer that generally do not overexpress HER2 and estrogen receptors. Expectedly, cancer cells of basal tumors had ∼60-fold and 250-fold lower expression of HER2 (*ERBB2*) and estrogen (*ESR1*) receptors as compared to cancer cells of HER2+ and Luminal subtypes, respectively (Suppl. Figure 12). NOTCH1 receptor expression was elevated in cancer cells of basal tumors, with *MFAP2* and *JAG1* as the top autocrine and paracrine specific ligands, respectively (Fig 5a-c, Suppl. Table 1). Consistent with this observation, cancer cells of basal tumors showed gene expression signatures of notch pathway activation relative to non-basal tumors (Suppl. Figure 13). Cancer cells of basal tumors also showed upregulated *FZD7* receptor expression and enrichment of autocrine *WNT11* and autocrine/paracrine *WNT3* interactions. Gene set enrichment analysis indicated an overall enrichment for activation of WNT/beta-catenin signaling in cancer cells of basal tumors (Suppl. Figure 13). Interestingly, the Frizzled/Wnt ligand antagonist, *SFRP1*, was inferred to have high expression in cancer cells of basal tumors while being nearly absent in cancer cells of other breast cancer subtypes (Fig 5c). At the receptor tyrosine kinase interface, cancer cells of basal tumors were characterized by increased *KIT* and decreased *RET* signaling compared to non-basal tumor types. Our analysis also inferred strong up-regulation of IL-6 signaling through GP130 (*IL6ST*) in cancer cells of non-basal tumors.

**Figure 5:**
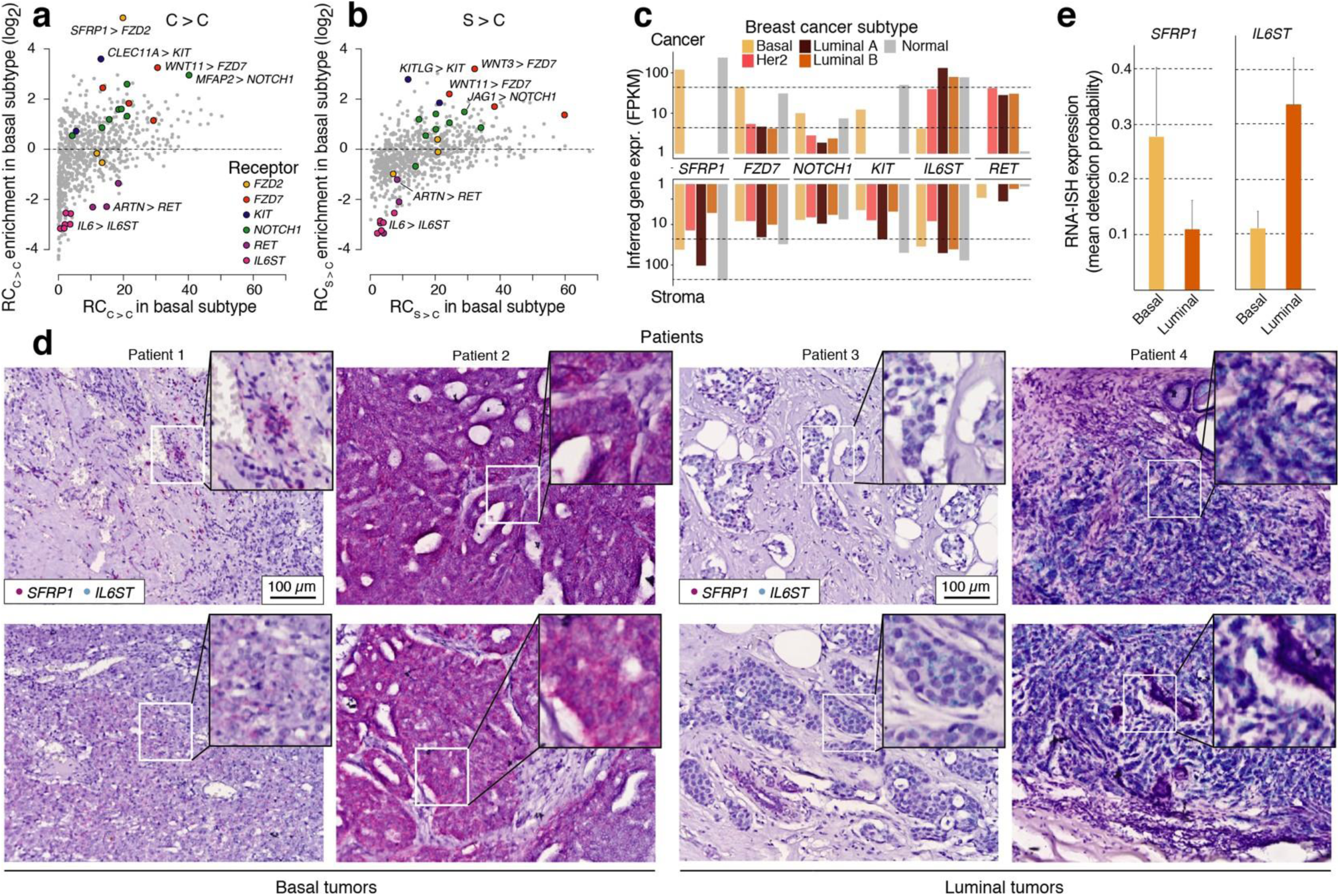
Inference of crosstalk in basal breast cancer. **a, b**) Enrichment of RC scores in breast cancer tumors of basal subtype for **a**) autocrine cancer-to-cancer and **b**) paracrine stroma-cancer crosstalk. **c**) Inferred expression of selected receptors and ligands in cancer and stromal compartment across breast cancer subtypes, expression in normal tissue (non-deconvolved) included for comparison. **d**) RNA ISH of *SFRP1* and *IL6ST* in 2 basal and 2 luminal FFPE breast tumor samples (2 slides per tumor). **e**) Quantification of *SFRP1* and *IL6ST* expression (mean detection probability across cancer cell regions) in the tissue sections using a supervised classification approach.

Next, we wanted to validate the TUMERIC predicted differential expression of *SFRP1* and *IL6ST* in the cancer cells of basal versus luminal breast cancer subtypes. We performed RNA in situ hybridization (ISH) using specific RNAScope probes generated against either *SFRP1* or *IL6ST* in FFPE sections of breast tumors from each subtype (see Methods, Fig 5d). Cancer cell mRNA expression in each subtype was quantified using two different approaches yielding similar results (Fig 5e and Suppl. Fig 14). Remarkably, and consistent with the TUMERIC prediction, we observed higher expression of *SFRP1* in the cancer cells of the basal subtype, compared to the luminal tumors (Fig 5e). In contrast, *IL6ST* had lower expression in cancer cells of the basal tumors as compared to the luminal subtype.

Overall, these data highlight putative differences in crosstalk of basal and non-basal breast cancer tumors and demonstrates how our approach can be applied to study cell signaling associated with specific molecular, genetic, or clinical subtypes of tumors.

## Discussion

Tumors are heterogenous mixtures of cancer cells and infiltrating non-cancer (stromal) cells. Molecular measurements obtained from a bulk tumor sample reflect an average across these components. By chance, the composition of these components will vary across tumor samples, introducing noise for downstream integrative analysis. However, in this study we demonstrate how this variability also enables inference of (average) cancer and stromal cell gene expression profiles across a collection of tumor samples. Our approach, TUMERIC, uses genomic data to first produce unbiased tumor purity estimates followed by constrained regression to deconvolve the matched bulk tumor transcriptome profiles. We show that this approach consistently recovers known hallmarks of cancer and stromal cell transcriptomes and is concordant with orthogonal single cell transcriptome and immunohistochemistry imaging data. The general methodology is in theory not restricted to transcriptomic data and might be used to deconvolve other types of bulk tumor molecular data such as proteomic (Fig 3e) or epigenetic profiles.

By applying the method to ∼8000 tumor samples across 20 solid tumors, we inferred ligand and receptor expression profiles in cancer and stromal cells across tumor types (Suppl. Table 1). We then nominated potential crosstalk within and between cancer and stromal compartments across tumor types. A large number of crosstalk interactions were inferred to be highly stroma specific across all tumor types. These interactions included many chemokine interactions as well as therapeutically important PD-1 and CTLA-4 immune checkpoint interactions. Checkpoint receptors were expectedly exclusively expressed in stromal cells. However, we inferred substantial expression of especially PD-L1 and CD86 ligands in the cancer cells of many tumor types, consistent with the hypothesis that cancer cells can express these ligands to attenuate T-cell responses (Sharma and Allison, 2015). Unexpectedly, most tumor types showed even higher expression of these checkpoint ligands in the stromal compartment, suggesting that the bulk of immune checkpoint inhibitory signals in the tumor microenvironment may be mediated by non-cancer cells. Interestingly, the tumor types with highest (e.g. lung squamous and kidney cancers) and lowest (e.g. colorectal cancer) inferred PD-L1 cancer cell expression have markedly different response rates to PD-L1/PD-1 checkpoint inhibition in clinical trials (Yarchoan et al., 2017). Further studies are however needed to explore the hypothesis that bulk tumor deconvolution of immune checkpoints can provide novel insights into the conditions for effective immune checkpoint therapy.

Our inferred recurrent cancer-cell autocrine LR pairs included the *MST1R* (RON) receptor, which acts upstream of the RAS-ERK and PI3K-AKT signaling pathways and is a prognostic marker and candidate therapeutic target in many solid cancer types (Yao et al., 2013). Additionally, LR interactions involving different low-density lipoprotein receptor-related proteins (LRPs) were frequently recurrently directed towards cancer cells (e.g. *WNT3*-*LRP6, APOE*-*LRP5, LTF*-*LRP11*), corroborating the importance of Wnt-signaling on cancer cell biology. Furthermore, our analysis showed that a recently proposed non-canonical *CXCR4* ligand, *CEL* (Panicot-Dubois et al., 2007), is frequently up-regulated in cancer cells of many tumor types, indicating a mechanism by which cancer cells might perturb *CXCR4* signaling in addition to the canonical *CXCL12*-*CXCR4* axis (Guo et al., 2015).

Cancer cell specific cross-talk interactions were often variable across tissues, underscoring the notion that cancer cell signaling is often tissue specific and likely dependent on the cell of origin. However, inferred cancer-cell autocrine cross talk often converged on Wnt, TGF-beta, Ephrin, and FGFR-family signaling pathways. The importance of these pathways in tumorigenesis is well established (Hanahan and Weinberg, 2011), underscoring the validity and utility of our approach. Moreover, the top cancer cell-specific interactions in individual tumor types highlight specific putative cancer cell signaling dependencies. Further functional studies are needed to validate these interactions and test whether they represent vulnerabilities that could be targeted therapeutically. Encouragingly, the inferred cancer cell specific interactions in brain tumors were highly concordant with previous studies. Firstly, the top inferred autocrine interaction was *DLL1*-*NOTCH1*, an autocrine/juxtacrine interaction critical for glioma tumorigenesis (Purow et al., 2005; Teodorczyk and Schmidt, 2015). Secondly, the top stroma-to-cancer specific interactions converged on *EGFR*, and all canonical *EGFR* ligands were overexpressed in the stroma. Stroma-to-cancer crosstalk is interesting because it comprises interactions and dependencies that are potentially lost when cancer cells are cultured *in vitro* or engrafted in a non-human microenvironment. Indeed, GBM cancer cells rapidly lose *EGFR* amplification and overexpression following *in vitro* cell culture (Pandita et al., 2004; William et al., 2017), however, amplification and overexpression can be maintained if the cancer cells are co-cultured with EGF ligands (William et al., 2017). Overall, these observations are consistent with our data and suggest that our methodology and data could help design *in vitro* assays and co-culture models that more accurately mimic the biology of the human TME.

Furthermore, we demonstrate that TUMERIC can nominate differences in crosstalk between molecular or clinical subtypes of a given tumor type. By inferring crosstalk specific to basal breast cancers, we recovered the expected patterns for HER2 and estrogen receptor expression across breast cancer subtypes. We inferred specific Notch and Wnt autocrine cancer cell signaling interactions specific to basal tumors. Interestingly, the Wnt inhibitory ligand, *SFRP1*, was inferred to have very high expression in cancer cells of basal tumors while being nearly absent in cancer cells of other breast cancer subtypes (Fig 4d). Downregulation of SFRP1 has been linked to epithelial to mesenchymal transition (EMT) in breast cancer cells (Scheel et al., 2011), indicating potential differences in EMT pathways or states between luminal and basal breast cancer cells. Interestingly, our analysis also inferred strong up-regulation of IL-6 signaling through GP130 (*IL6ST*) in cancer cells of non-basal tumors, consistent with previous studies highlighting a functional role for RET-IL6 crosstalk in ER-positive breast tumors (Gattelli et al., 2013). Importantly, we could validate the predicted subtype specific expression of *SFRP1* and *IL6ST* (GP130) using RNA-ISH in breast cancer FFPE tissue sections. Overall, these data suggest that tumor subtypes can have profound differences in crosstalk, and our analysis can provide a list of putative, testable interactions underlying breast cancer subtypes.

We note that a key limitation of our approach is that local concentrations of ligands and receptors may differ considerably from our average compartment-level estimates. While this is especially important when considering between-compartment crosstalk, it likely matters less for the inference of cancer-to-cancer autocrine signaling (where the same cell produces the interacting ligand and receptor). We expect future high-throughput technologies such as sample-level spatial transcriptomics could be combined with our cohort-scale approach to provide a more accurate and comprehensive description of tumor crosstalk.

Overall, our study provides unique insights into the complex interactions shaping the TME. Our approach is especially useful in settings where bulk tumor biopsy data is either already abundant or the only feasible data source, which is the case in many clinical settings. We anticipate that the method could complement existing biomarker and target discovery approaches by providing computationally purified molecular profiles of cancer cells as they exist inside human tumors.

## Methods

### Tumor data sources

We analyzed 20 solid tumor types with TCGA acronyms BLCA, BRCA, CESC, CRC (COAD and READ combined), ESCA, GBM, HNSC, KIRC, KIRP, LGG, LIHC, LUAD, LUSC, OV, PAAD, PRAD, SKCM, STAD, THCA and UCEC. The full names of cancer types and sample size are indicated in the Suppl. Table 1. We obtained somatic mutation (SNV) and copy number variation (CNV) data for 20 tumor types from the Broad Institute Firehose website (See data availability section below). Uniformly processed TCGA RNA-seq (FPKM) data was obtained from the UCSC Xena server.

### Tumor purity estimation

We used 4 different published methods for consensus tumor purity estimation: AbsCNseq, PurBayes, Ascat and ESTIMATE. AbsCNseq uses CNA segmentation and SNV variant allele frequency (VAF) data of individual tumors(Bao et al.); PurBayes utilizes SNV VAF data of diploid genes (inferred from CNA data)(Larson and Fridley); Ascat purity estimation is based on CNA (SNP array) data, where tumor ploidy and purity are co-estimated to identify allele specific CNA(Loo et al.), pre-computed Ascat tumor purity estimates for the TCGA cohort were obtained from the COSMIC website (See data availability section). ESTIMATE uses mRNA expression signatures of known immune and stromal gene signatures to infer tumor purity(Yoshihara et al., 2013), and tumor purity values were obtained by applying ESTIMATE to the TCGA RNA-seq (log2 FPKM) data. In order to derive consensus tumor purity estimates, we carried out missing data imputation followed by quantile normalization separately for each cancer type. Some tumor purity values were missing because the algorithms failed to converge on certain input data. Additionally, we observed some instances of very high (>98%) or low (<10%) purity estimates, but such cases were usually only found by a single method for a given tumor and were therefore also assigned as missing data. Missing data was then imputed using an iterative Principal Component Analysis of the incomplete algorithm-vs-sample tumor purity matrix (using the missMDA R package(Josse and Husson, 2016)). We used quantile normalization to further align and standardize the tumor purity distributions of different algorithms per tumor type. Briefly, a mean reference purity distribution is computed and these mean values are substituted back into the purity distributions of the individual algorithms. Since ESTIMATE generated purity estimates with a large bias compared to the other three methods (generally 30-50% higher, Suppl. Fig 1), we did not use ESTIMATE purity values to compute the mean reference distribution (see code in Supplementary Data). The final TUMERIC consensus tumor purity estimate was obtained as the mean of these normalized purity values.

### Comparison of transcriptome deconvolution methods

TCGA RNA-seq HTSeq count data for BRCA, GBM, HNSC, LGG, LUAD, LUSC, SKCM and THCA tumor types (used for DeMixT (Wang et al., 2018), LinSeed (Zaitsev et al., 2019)) and TPM data for HNSC, LUAD, LUSC tumor types (used for CiberSortX (Newman et al., 2019)) were downloaded from the UCSC Xenahub platform (https://xenabrowser.net/). All methods were executed on 100 randomly selected tumor transcriptome samples from each tumor type. The datasets were normalized and parameters were set according to authors instructions (see Supplementary Methods). CiberSortX could only be run for tumor types where a compatible signature matrix was available (HNSC, LUAD, and LUSC), and DeMixT could only be run for tumor types with available normal samples (all selected tumor types except SKCM). DeMixT and LinSeed were run using R v.3.6.1 and their respective R packages, CiberSortX was run using its online web interface. Estimated tumor purity fractions were compared to values computed by the TCGA pan-cancer consortium using the ABSOLUTE algorithm (https://gdc.cancer.gov/about-data/publications/pancanatlas).

### Database of ligand-receptor interactions

We obtained ∼1400 ligand-receptors pairs supported by evidence in the literature as compiled and curated by Ramilowski et al. (Ramilowski et al., 2015). 7 additional known immune checkpoint interactions (Khalil et al., 2016) were manually added to the analysis: CD274-PDCD1, CD80-CTLA4, CD80-CD28, NECTIN2-CD226, NECTIN2-TIGIT, PVR-TIGIT, SIGLEC1-SPN. The complete list of ligand-receptor interactions can be found in Suppl. Table 2.

### Cancer-stroma gene expression deconvolution

We assume tumors to be comprised of cancer and stromal (any non-cancer) cells. Measured bulk tumor mRNA abundance is then given by the sum of mRNA molecules derived from these two compartments. Tumor mRNA expression (*e*_”#$%&,(_) measured for a given gene in sample *i* can then be expressed as:

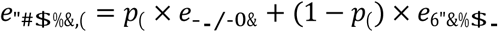

Here *p*_*i*_ denotes the cancer cell proportion (tumor purity) in sample *i*, and *e*_-./-0&_ and *e*_6”&%$._ are average expression levels for the gene in the cancer and stromal compartment (not dependent on sample *i*), respectively. We make the simplifying assumption that these (non-negative) average compartment expression levels are constant across the set of tumors and estimate them using non-negative least squares regression. 95%-confidence intervals and standard deviations for the cancer and stroma point estimates are estimated using bootstrapping.

Bulk tumor RNA-seq FPKM data was log-transformed, log_2_(X+1), before deconvolution. It has previously been discussed whether mixed-tissue gene expression deconvolution should be done using linear or log-transformed gene expression values(Shen-Orr et al., 2012; Zhong and Liu, 2012). Firstly, we observed that the relationship between tumor purity and raw bulk tumor gene expression was generally heteroscedastic: Across all cancer types, tumor purity had overall stronger linear correlation with log-transformed RNA-seq gene expression data, as can be observed in deconvolution of known stromal-specific genes (Suppl. Fig 2-5). Secondly, we directly compared results for raw and log-transformed data, and while the results were overall similar, we found that log-transformation provided better separation between inferred cancer and stroma compartment gene expression for known stromal genes (Suppl. Fig 15), which is consistent with previous empirical analysis (Shen-Orr et al., 2012).

We found that the equation above tended to misestimate stromal gene expression for genes with somatic copy number alterations (CNA) affecting gene expression in a subset of the samples (for example *ERBB2* in HER2-positive breast tumors). We therefore used a modified approach for such genes. We first identified genes with correlation between CNA and mRNA expression (comparing expression for samples with diploid and non-diploid CNA, Mann-Whitney U-test, P<1e-6, to account for multiple testing) in a given set of tumors, and then estimated cancer and stromal compartment gene expression using a two-step approach. Stromal compartment mRNA expression was first inferred using the above approach using only samples with diploid copy number for the gene. We then used the inferred mean stroma compartment expression, the measured mean tumor expression, and the mean purity of the tumor samples to calculate the mean cancer compartment expression using the above equation.

A limitation of TUMERIC is that the method estimates average cancer and stromal compartment gene expression across a cohort of tumor samples. Since the stromal compartment is a mix of many different types of cells, the averaged stromal expression estimates therefore require careful interpretation. Furthermore, exploration of biologically relevant between-sample variation in gene expression, not due to variation in tumor purity, requires additional downstream analysis of the expression residuals.

### Deconvolution of iTRAQ tumor protein abundance data

We obtained iTRAQ protein abundance data for BRCA and OV tumor types using CPTAC consortium data available at cBioPortal (www.cbioportal.org). Bulk abundance profiles were deconvolved into cancer and stroma compartment abundance profiles similar to the procedures described for RNA-seq data above. Briefly, we have matched protein abundance and consensus tumor purity data available for each tumor. We can therefore use the consensus tumor purity estimates and NNLS to infer the (mean) abundance of individual proteins in the cancer and stromal compartment, respectively.

### Ligand-Receptor Relative Crosstalk (RC) score

Following the law of mass action, we can estimate the concentration of a ligand-receptor (LR) complex from the molar concentrations of the individual ligand, *L*, receptor, *R*, and their dissociation constant, *K*_*D*_, under equilibrium:

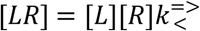

In our analysis we are only evaluating relative changes of individual LR complex concentrations across samples or compartments. Since the dissociation constant cancels out (present in numerator and denominator) when we compute relative concentrations (RC score below) we can ignore the dissociation constant in downstream analysis. Our analysis depends on multiple simplifying assumptions. We assume that compartment-level mRNA expression estimates are reasonable proxies for ligand and receptor molar concentration at the site of LR-complex formation. Additionally, we assume there is no external competition for individual ligands and receptors of a given LR pair, or that these effects are constant across samples or compartments.

To estimate the relative flow of signaling between cancer and stromal cell compartments, we developed the Relative Crosstalk (RC) score. LR complex activity is first approximated using the product of ligand and receptor gene expression inferred for the given compartments (in linear scale). The RC score then estimates the relative complex concentration given all four possible directions of signaling and a normal tissue state, e.g. for cancer-cancer (CC) signaling:

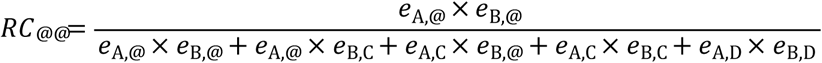

The normal term in the denominator is included to normalize for complex activity in normal tissue, and this term is calculated directly from the observed gene expression levels in normal tissue samples available for each tumor type in TCGA.

### Gene-set enrichment analysis

To study genes differentially expressed between cancer and stromal compartments, we performed GSEA (Subramanian et al., 2005) pre-ranked analysis of genes sorted by differential expression (log fold-change) in the two compartments. We analyzed all hallmark gene sets and used a FDR cut-off of 0.25 to determine gene sets with differential enrichment.

### Immunohistochemistry (IHC) quantification analysis of Human Protein Atlas data

In order to quantify cancer and stromal cells expression of genes, we performed color deconvolution of IHC images obtained from the Human Protein Atlas (proteinatlas.org) (Uhlen et al., 2017) using the ImageJ software package and standard protocols (Schindelin et al., 2015). Following manual selection and segmentation of cancer and stromal cells (without knowledge of antibody staining), color intensities were measured with ImageJ, and DAB (target), hematoxylin (cells), and complementary components were estimated. Average antibody intensities were then estimated for the cancer and stromal compartment of a given slide. Further technical details are provided in the supplementary methods.

### RNA-ISH of breast cancer tissue

We obtained Formalin-Fixed Paraffin-Embedded (FFPE) breast cancer tissue slides for 2 luminal (ER+/PR+) and 2 triple negative basal-like breast tumors. In situ hybridization (ISH) was performed using specific RNAScope (https://acdbio.com) probes to evaluate IL6ST and SFRP1 RNA level and localization. The FFPE slides were deparaffinised, rehydrated, and pretreated using the RNAScope Sample Preparation kit according to the manufacturer’s recommendations. The slides were incubated in target retrieval solution (#322000, Advanced Cell Diagnostics) for 15 minutes at 97°C followed by a protease solution (#322330, Advanced Cell Diagnostics) for 30 minutes at 40°C. RNAScope IL6ST (#447251) and SFRP1 (#429381-C2) target probe and RNAScope 2.5 HD Duplex Detection (Chromogenic) kit (#322430, Advanced Cell Diagnostics) were applied to the slide according to the manufacturer’s instructions. RNA expression was quantified using two approaches. Firstly, according to the manufacturer’s instructions (TS 46-003), we segmented the Images using supervised machine learning. Briefly, we used Fiji/ImageJ to train a 4-class random forest classifier to distinguish regions with *SFRP1* staining, *IL6ST* staining, nuclear regions, and background signal. We used this model to segment all 8 images. We selected 5 cancer cell regions in each image and recorded the mean probability for the *SFRP1* and *ILST6* staining classes in each region. These staining probabilities were summarized (mean and standard deviation) across all regions from the four images (2 patients) for the luminal and basal subtype, respectively. As a second and different approach, we quantified *IL6ST* and *SFRP1* mRNA expression in cancer cells of basal and luminal breast cancer tissues using 3-channel color deconvolution. Channels were specified for probe 1 (*SFRP1*), probe 2 (*IL6ST*), and background. Multiple areas enriched for cancer cells were selected in each tissue slide and the optical density (OD) was calculated for each area. The average and standard deviation of the OD for each gene and tissue type was calculated.

## Supplementary data

All code used to generate the figures and statistics of the paper is included in Supplementary Data 1. Source code for Tumeric is attached as Supplementary data 2. RC scores and other summary metrics for all analyzed ligand-receptor pairs are summarized in Supplementary Table 1. Supplementary data files can be accessed at: https://bit.ly/2Ingp03.

## Data availability

TCGA SNV and CNV data: https://gdac.broadinstitute.org/. Data release version 2016/01/28. TCGA RNA-seq data: https://toil.xenahubs.net/download/tcga_RSEM_gene_fpkm.gz ASCAT purity estimates: https://cancer.sanger.ac.uk/cosmic/download

IHC data: https://www.proteinatlas.org.

## Author contributions

AJS conceived of the project. UG, MMN, NR, TTN, and AJS analyzed data. UG, SS, and AJS designed and implemented methods and software. JY, PHT, JI and BC acquired and annotated FFPE samples. AW, RD, and AJS performed RNAscope analysis. AJS, UG, and RD wrote the manuscript.

## Acknowledgements

This research is supported by the Singapore Ministry of Health’s National Medical Research Council under its OF-IRG program (OFIRG18may-0075), and Agency for Science, Technology and Research (A*STAR) under its CDAP program (grant no. 1727600057).

